# Computational Screening of Potential Inhibitors of *Desulfobacter postgatei* for Pyrite Scale Prevention in Oil and Gas Wells

**DOI:** 10.1101/327957

**Authors:** Abdulmujeeb T. Onawole, Ibnelwaleed A. Husseinl, Mohammed A. Saad, Musa E.M. Ahmed, Hassan I. Nimir

## Abstract

Sulfate-reducing bacteria (SRB) such as *Desulfobacter postgatei* are often found in oil and gas wells. However, they lead to the release of hydrogen sulfide which in turn leads to the formation of iron sulfide scale such as pyrite. ATP sulfurylase is an enzyme present in SRB, which catalyzes the formation of adenylyl sulfate (APS) and inorganic pyrophosphatase (PPi) from ATP and sulfate which is one of the first steps in hydrogen sulfide production by *D. postgatei*. Virtual screening using molecular docking and machine learning tools was used to identify three potential inhibitors of ATP sulfurylase from a database of about 40 million compounds. These selected hits ((S,E)-1-(4-methoxyphenyl)-3-(9-((m-tolylimino)methyl)-9,10-dihydroanthracen-9-yl)pyrrolidine-2,5-dione;,methyl 2-[[(1S)-5-cyano-2-imino-1-(4-phenylthiazol-2-yl)-3-azaspiro[5.5]undec-4-en-4-yl]sulfanyl]acetate and (4S)-4-(3-chloro-4-hydroxy-phenyl)-1-(6-hydroxypyridazin-3-yl)-3-methyl-4,5-dihydropyrazolo[3,4-b]pyridin-6-ol), which are known as A, B and C respectively) all had good binding affinities with ATP sulfurylase and were further analyzed for their toxicological properties. The molecular docking results showed that all the compounds have negative binding energy with compound A having the highest docking score. However, based on the physicochemical and toxicological properties, compound C is the best choice as it does not violate any of the recommended properties that relate to absorption and distribution. Only compound C was predicted to be both safe and effective as a potential inhibitor of ATP sulfurylase. The binding mode of compound C revealed favorable interactions with the amino residues LEU 213, ASP 308, ARG 307, TRP 347, LEU 224, GLN 212, MET211 and HIS 309.

**Importance:** Scale formation formed by hydrogen sulfide, which is produced by sulfate reducing bacteria such as *Desulfobacter postgatei* has been a persistent problem in the oil and gas industry leading to loss of money, time and even lives. The three selected hits from the virtual screenings of about 40 million compounds would possibly inhibit the enzyme, ATP sulfurylase, which is involved in the first reaction in hydrogen sulfide formation in *Desulfobacter postgatei*. The selected inhibitors are expected to significantly reduce the formation of hydrogen sulfide and consequently prevent the development of pyrite scale in oil and gas wells.

## 1. Introduction

Sulfate-reducing bacteria (SRB) have been a persistent problem in the oil and gas industry as their presence in oil and gas reservoirs is abundant. SRB are one of the main sources of hydrocarbons as one of the predominant sources of sulfide and sulfur during the maturation of oil reservoirs (1, 2). They help in sulfur formation by the incomplete oxidation of sulfur. This sulfur in turn leads to serious problems such as iron sulfide (pyrite) scale formation which hinders the injectivity of water injection wells, corrosion of iron and contamination of produced gas by generating hydrogen sulfide (H_2_S) (3). These problems sometimes lead to deaths during oil and gas production such as the fatalities caused by hydrogen sulfide poisoning during offshore operations in the North Sea (4). SRBs have been tagged has the major perpetrators of microbially influenced corrosion (5, 6).

SRBs operate mostly in anaerobic conditions and use sulfate as an electron acceptor during their metabolism of energy (3). They are prevalent in water and land environments as long as they are anaerobic and such anaerobic environments includes oil and natural gas wells (7). *Desulfobacter postgatei* is one of the common SRBs that is rod like or elliptical in shape. Unlike other SRBs, *D. postgatei* converts acetate to carbon dioxide (CO_2_) and H_2_S (3, 8) as described in equation 1 below.

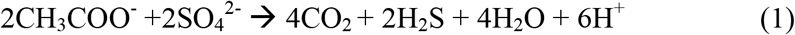

However, in the reduction of sulfate to H_2_S some enzymes are involved (Table 1) which make the reaction feasible. The first of these reactions is catalyzed by ATP sulfurylase (9, 10).

**Table 1.** Reactions involved in respiratory sulfate reduction

| Reaction | Enzyme Involved |
| --- | --- |
| $\text{SO}_4^{2-} + \text{ATP}^{4-} \rightarrow \text{APS}^{2-} + \text{PPi}^{4-}$ | ATP sulfurylase |
| $\text{PPi} + \text{H}_2\text{O} \rightarrow 2 \text{Pi}$ | Inorganic pyrophosphatase |
| $\text{APS} + \text{H}^+ + 2\text{e}^- \rightarrow \text{HSO}_3^- + \text{AMP}$ | APS reductase |
| $\text{HSO}_3^- + 6\text{H}^+ + 6\text{e}^- \rightarrow \text{HS}^- + 3\text{H}_2\text{O}$ | Bisulfite reductase |

Existing solutions to alleviate the effects of SRBs include the use of biocides like aldehydes and amines, which are often hazardous and mutagenic (11). In addition, dissolved oxygen and nitrates create an aerobic environment. however, a few SRBs thrive under aerobic conditions (7).Also, heavy metals are used but they result in problems of toxicity.

Recent works (12, 13) have proffered green formulations that can be used in scale removal. However, a root-cause analysis in preventing scale formation is likely the inhibition of the enzymes responsible for sulfate reduction present in SRB. Computational techniques such as molecular docking and machine learning have been extensively used in the pharmaceutical industry in discovering and designing novel compounds which inhibit certain proteins to create a pharmacological effect (14, 15). This process can be translated to oilfield chemistry in finding a safer and longer lasting solution to sulfur production by SRB. Moreover, these methods can help in screening large databases of compounds and also predict their toxicity properties which can help as a guide in selecting safe compounds that can be used as inhibitors of ATP sulfurylase (16, 17).

There is a dearth of literature using this approach in reducing sulfur production in oilfields. A relatively recent work (18) screened about 15 compounds to find potential inhibitors for *Archaeoglobus fulgidus*, which though not a bacterium but it is also sulfate reducing organism. In this work, we use computational techniques such as molecular docking and machine learning to select hits by screening from a database of about 40 million compounds and use homology modeling to model the target protein from the protein sequence of *D. postgatei*. The selected hits from this work have the potential to inhibit ATP sulfurylase, hence alleviating the threat of SRBs including pyrite scale formation in the oil and gas industry.

## 2. Methodology

### 2.1. Homology Modeling

Selecting a protein target is the first and most crucial step in structure based virtual screening. Unfortunately, as there was no structure of ATP sulfurylase in the protein databank (PDB) hence, the structure was designed using homology modeling with the aid of the Raptor program (19, 20). The protein sequence for the ATP sulfurylase enzyme (Fig. 1) was obtained from the genome portal of the Department of Energy Joint Genome Institute (21, 22). As homology models are dependent on the quality of an existing crystal structure. The model was based on the template of the crystal structure of ATP sulfurylase of *Allochromatium vinosum* (PDB ID: 4DNX) (23, 24) *with* a resolution of 1.6 Å. This implies that the resolution is high as electron density maps derived from the X-ray crystallographic data of a protein structure with a resolution of at most 2.0 Å are deemed high resolution structures (25). The structure was validated using a Ramachandran plot. Since the structure is not an X-ray crystal structure, which are often co-crystallized with ligands that indicate the binding site, hence blind docking process was employed. This process involves considering the whole of the protein during the molecular docking process to find potential binding sites and is used when a binding site is unknown (26–28). The PyRx (29) software was used to determine the binding parameters using the maximize option, which enclosed the entire protein molecule, since the binding site is not known and blind docking is being employed. This resulted in a binding site center of 2.0027, −1.9107 and 24.2677 for the X, Y and Z axes respectively. The size of the box for the docking was 100 × 100 × 100 Å^3^. These parameters ensured that the whole structure was covered during the virtual screening process.

**Figure 1.**
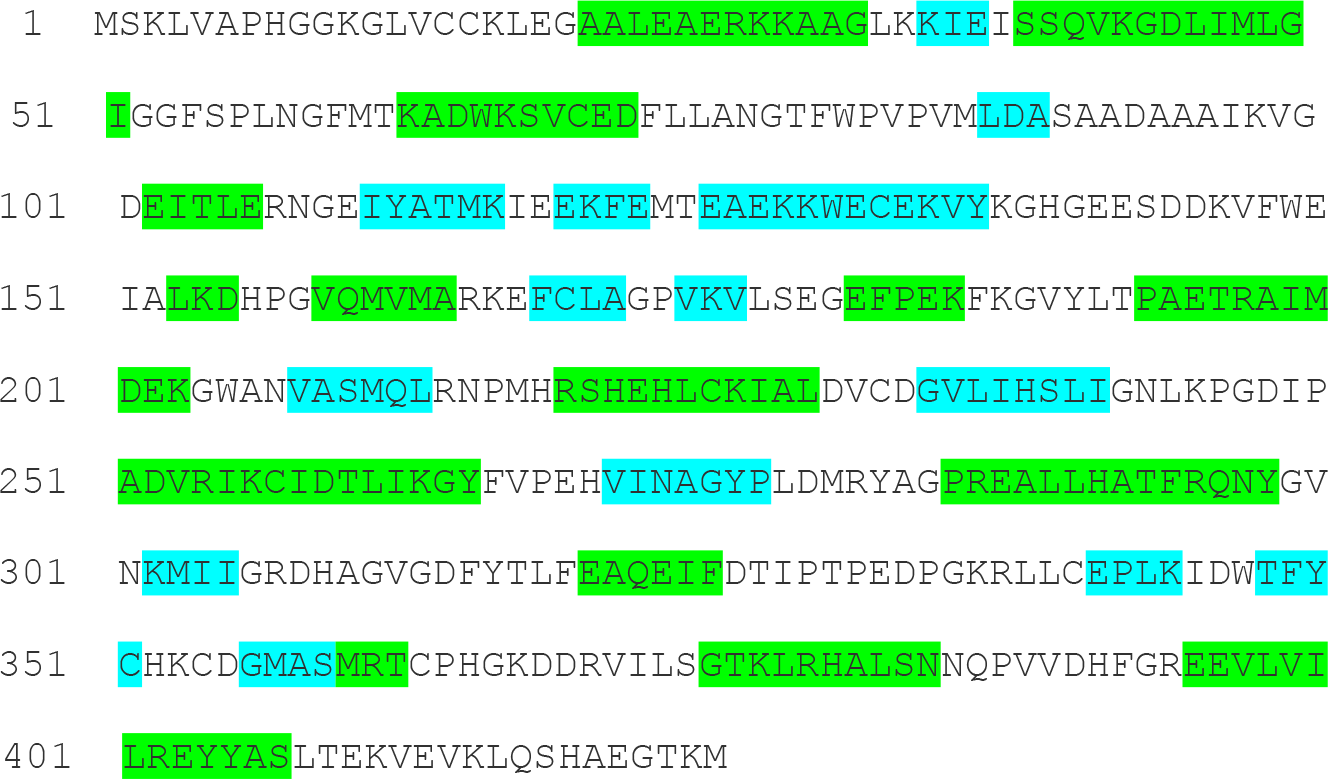
The protein sequence of ATP sulfurylase (*Desulfobacter postgatei*) in clustal format. (Green highlight-helix, Blue highlight-Strand, white-coil)

**Figure 2.**
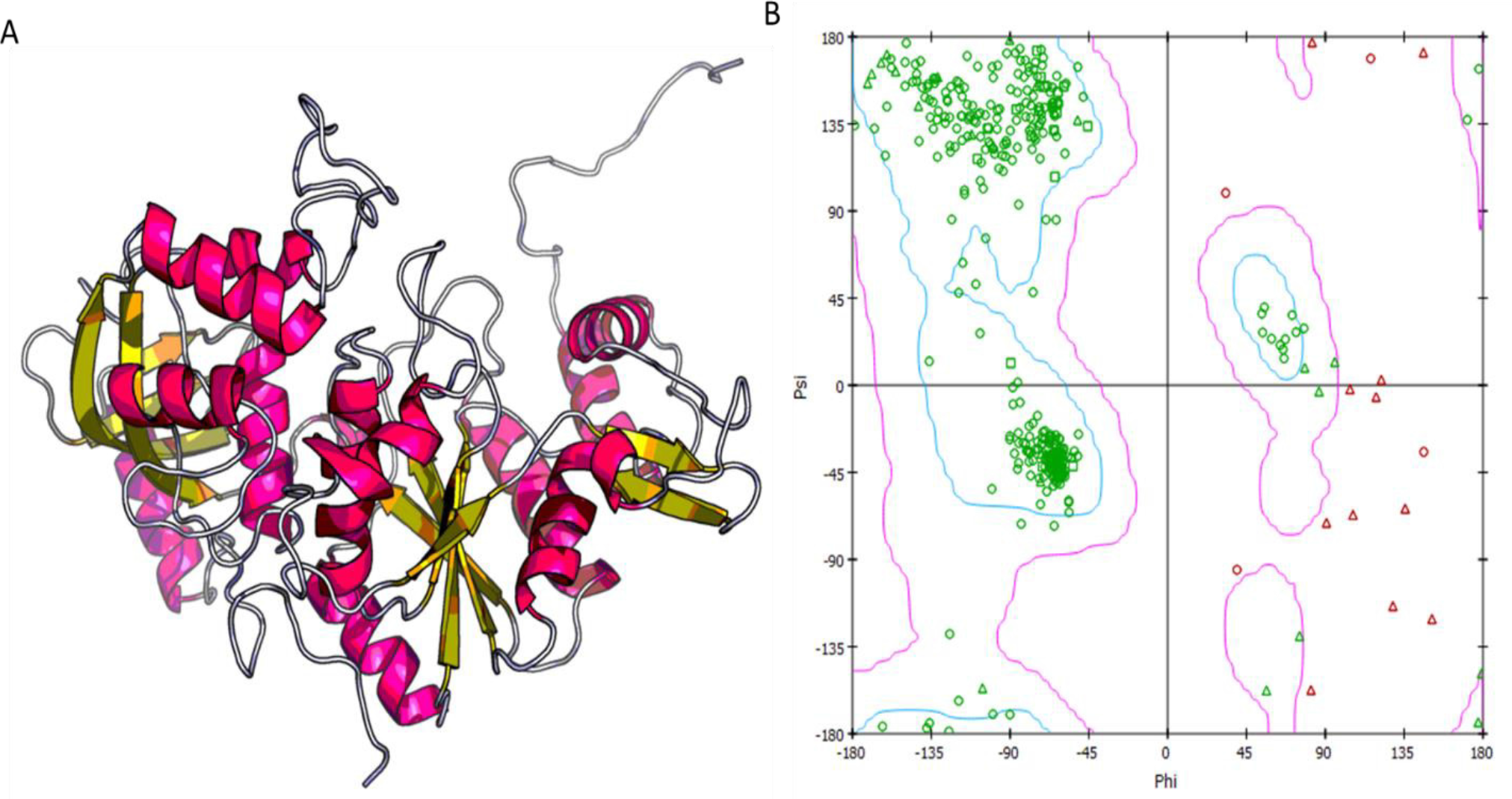
The (a) the tertiary structure and (b) Ramachandran plot of ATP sulfurylase (*Desulfobacter postgatei*) (Red-helix, yellow-Strand, white-coil;, squares - proline, glycine - triangles, squares - proline, others-circle).

### 2.2 Virtual Screening

The homology modeled structure was virtually screened against 100,000 compounds randomly chosen from the MCULE database (30), which consists of precisely 39,884,964 compounds at the time of this work.. To select the 100,000 compounds randomly, the basic properties option was used. This option filters compounds from the database on the basis of the following conditions: the filtered compounds should not violate more than one of the Lipinski’s rule of 5 (RO5) (31–33), the compounds should not exceed a maximum of 10 rotatable bonds, a maximum of 5 halogen atoms, a maximum of 5 chiral centers; a minimum of 10 heavy atoms and a minimum of 1 aromatic ring (15). AutoDock Vina was used for the molecular docking of the filtered compounds with the protein (34). This criterion of filtering aims at ensuring that the selected compounds had the desirable physicochemical properties, which are important during absorption and toxicity studies. Only the top scored 500 compounds were kept from the virtual screening.

### 2.3 Consensus Scoring

Consensus scoring is a process of combining of two or more scoring algorithms to rank the binding affinities of ligands to a target protein. Consensus scoring has been known to reduce the rate of false positives (35–37) while also serving as a validation tool for the binding affinities. Hence, the use of consensus scoring has led to improved hit rates. Machine learning has proved to be useful tool in calculating binding affinities in a much shorter time than molecular docking (38–40). The K_DEEP_ machine-learning tool, which uses Convolutional Neural Networks (CNN), was used for a second virtual screening. The K_DEEP_ gives the binding affinity result in pK_d_ such that the higher the pK_d_ value the stronger the binding affinity. The results of both virtual screenings (molecular docking and K_DEEP_) were normalized such that values close to 1 and 0 corresponded to the top and low scored values, respectively. A vote rank method was used in the consensus scoring process to select the hits which mutually appeared in the top 5% scored ligands from the two virtual screenings (14, 41).

## 3. Results and Discussion

### 3.1 Protein Structure and Validation

The homology modeled structure (Fig. 2a) depicted about 10 helices, which are connected by strands and coils. With the aid of the Ramachandran plot as a validation tool (Fig. 2b). The Ramachandran plot depicts the allowed values of ψ (Psi) versus φ (Phi) angles corresponding to many confirmations, for a specific amino acid (42). The structure is made up of 423 residues with 389, 27 and 7 being in the favored, allowed and outlier regions, respectively. The upper and lower left regions of the Ramachandran plot correspond to the β-pleated and right handed α-helices while the middle of the right region indicates the left-handed α-helices. For a superior quality protein, it is expected that the outlier percentage should not exceed 5% while the favored and allowed region should have a combined total of more than 90 % (43–45). The seven residues in the outlier make up 1.7 % while the favored and allowed region both constitute 98.3 %, hence the structure is of excellent quality and hence is reliable for molecular docking.

**Figure 3.**
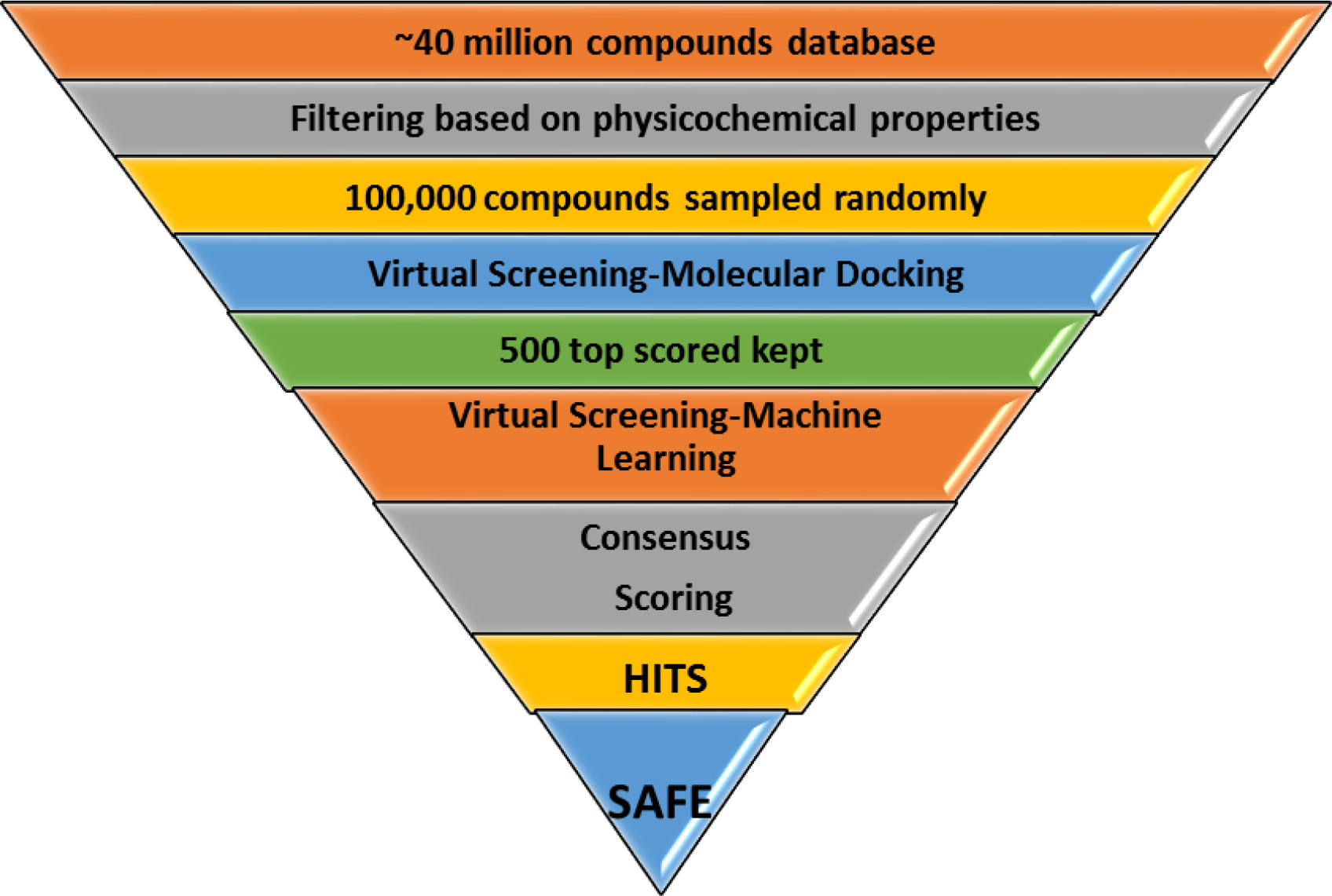
Methodology workflow of hit selection

### 3.2 Docking Simulations

Using the vote rank approach, the consensus scoring (Figs 4 and 5) from the two virtual screenings (molecular docking and machine learning) resulted in the selection of three compounds as hits namely: MCULE-798447760801 (A), MCULE-123221368002 (B) and MCULE-938606270701 (C), which have the following IUPAC names: (S,E)-1-(4-methoxyphenyl)-3-(9-((m-tolylimino)methyl)-9,10-dihydroanthracen-9-yl)pyrrolidine-2,5-dione; methyl 2-[[(1S)-5-cyano-2-imino-1-(4-phenylthiazol-2-yl)-3-azaspiro[5.5]undec-4-en-4-yl]sulfanyl]acetate and (4S)-4-(3-chloro-4-hydroxy-phenyl)-1-(6-hydroxypyridazin-3-yl)-3-methyl-4,5-dihydropyrazolo[3,4-b]pyridin-6-ol. These compounds would henceforth be referred to as compounds A, B and C (Fig. 6). Compound A was the 5^th^ and 21^st^ top scored compound for Auto Dock Vina and K_DEEP_ respectively, while compound B correspond to the 11^th^ and 3^rd^ top scored compound for Auto Dock Vina and K_DEEP_ respectively. Compound C was the 17^th^ and 20^th^ top scored compound for Auto Dock Vina and K_DEEP_ respectively. All the selected compounds had a negative free energy of binding from the molecular docking score which corresponds to −8.7, −8.4 and −8.2 kcal/mol for compounds A, B and C, respectively. The negative values of the docking score implied that they all bound to the target protein with compound A having the highest binding energy. The pK_d_ values from K_DEEP_ were 6.56, 7.03 and 6.58 for compounds A, B and C respectively with compound B having the highest pK_d_ value (Tables S1 and S2-Supproting information).

**Figure 4.**
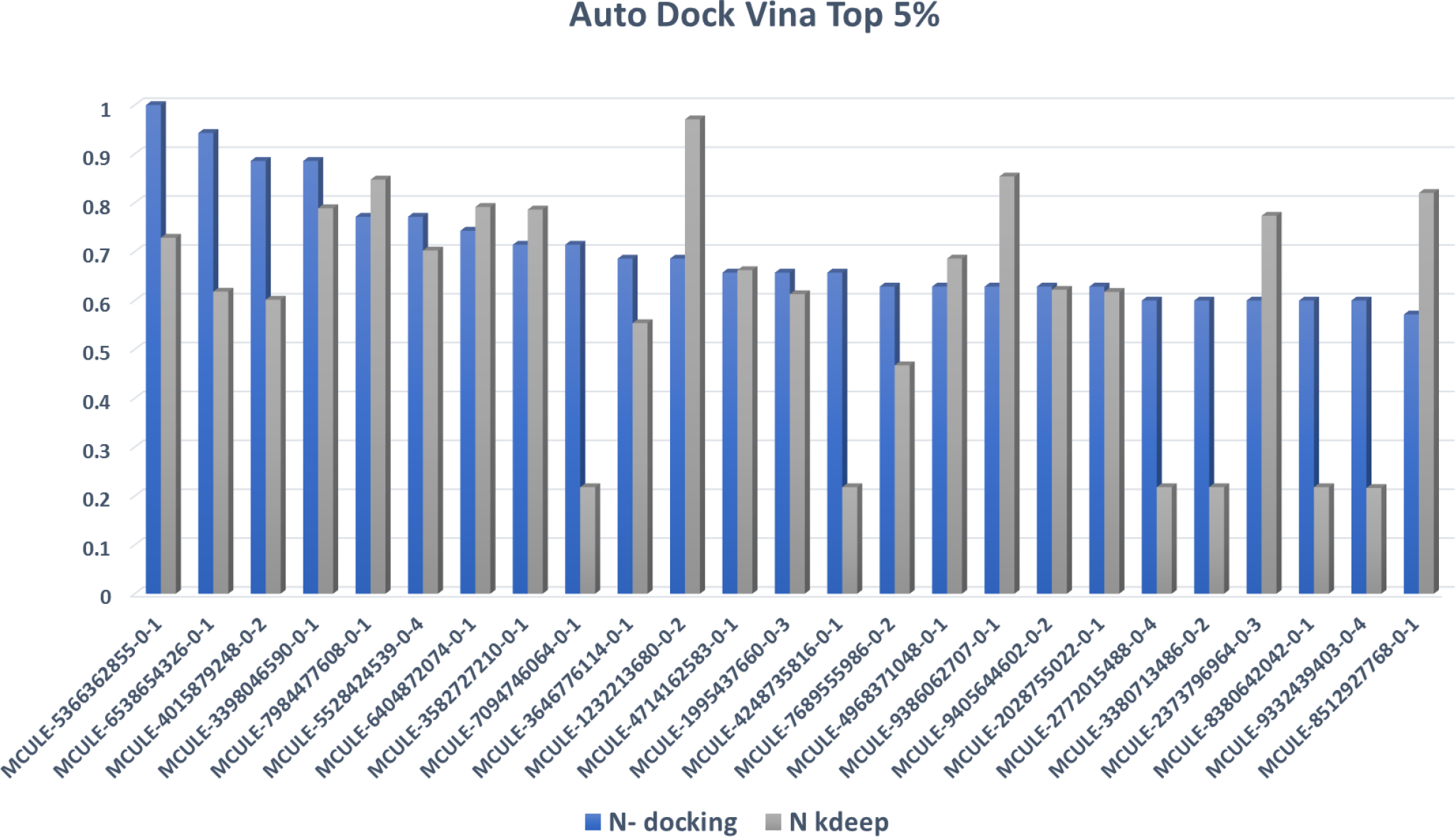
Top 5% normalized scored compounds from Auto Dock Vina (Molecular docking)

**Figure 5.**
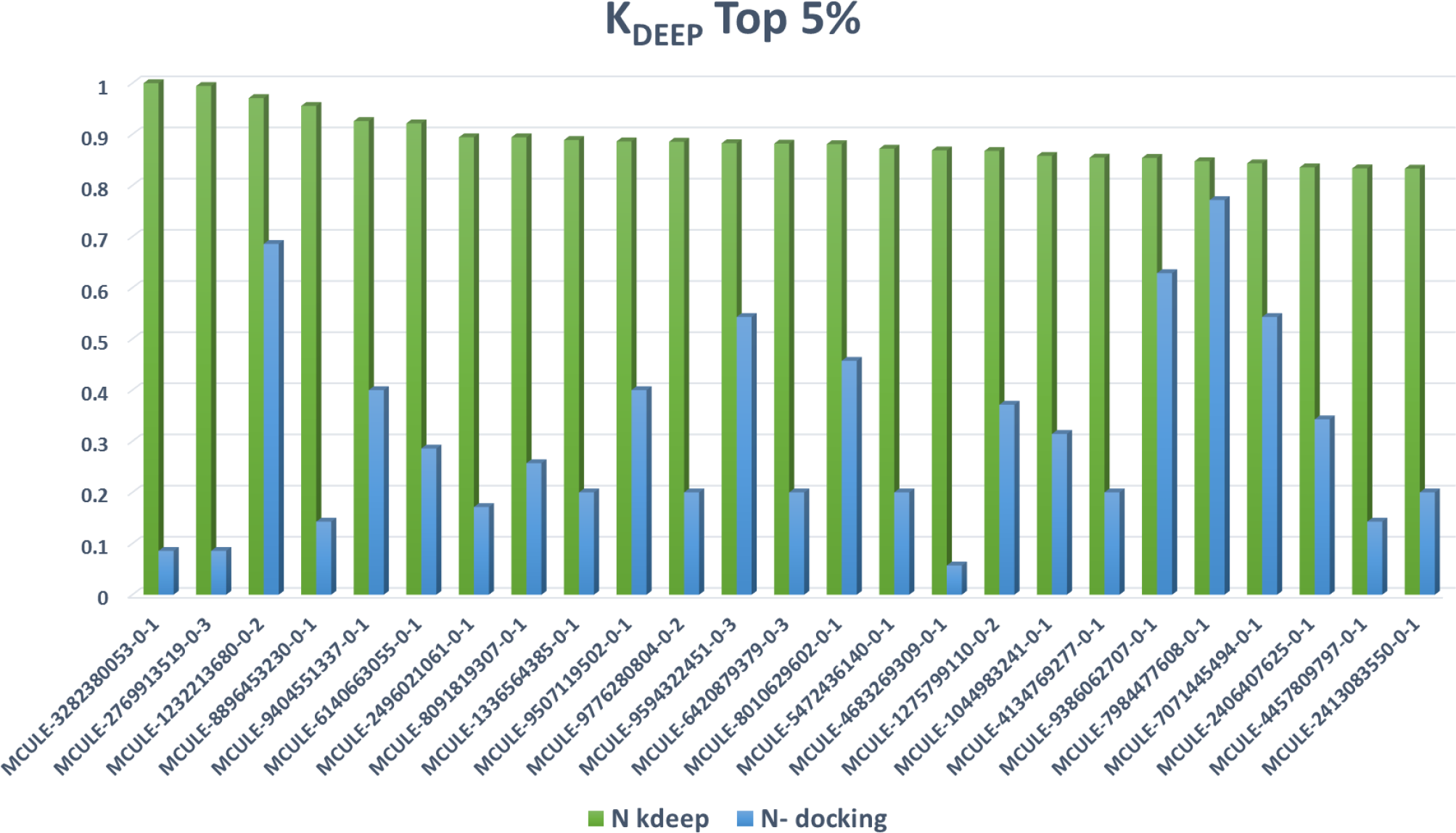
Top 5% normalized scored compounds from K_DEEP_ (Machine Learning)

**Figure 6.**
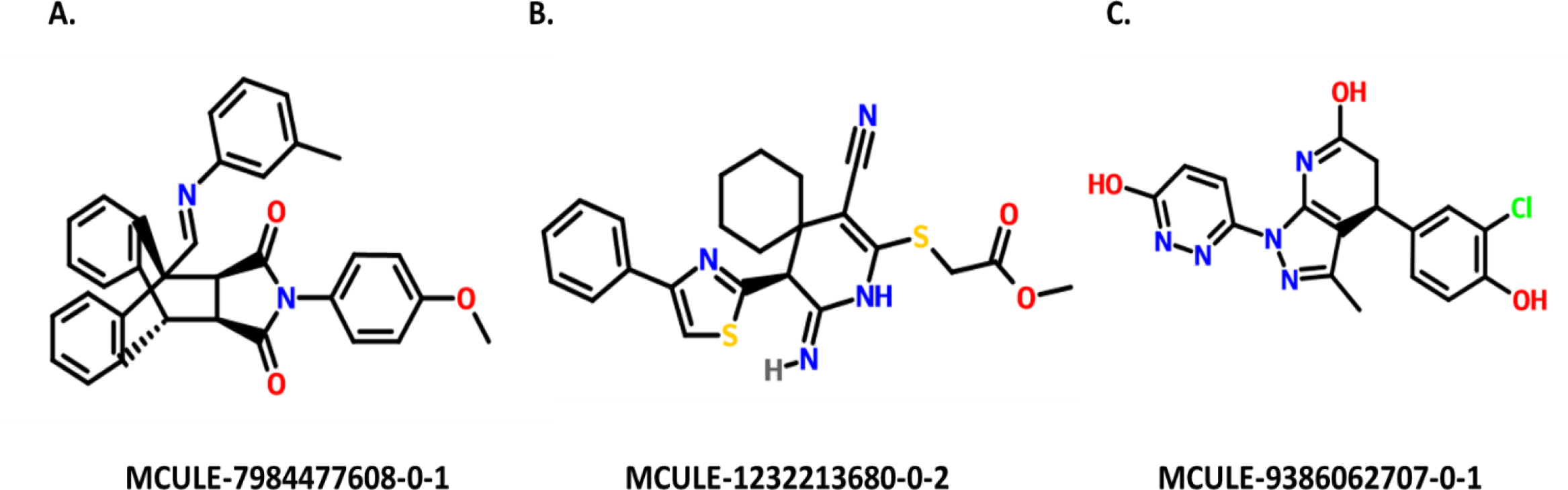
Selected compounds as HITS from consensus scoring

### 3.3 Binding Mode and Molecular Interactions of Selected Compounds

The molecular docking provided insight into the mode of binding and the molecular interactions of the selected compounds in the protein target. Compound A showed many favorable molecular interactions with the amino residues in the binding pocket of the target protein (Fig. 7). They include hydrogen bonding with GLU 287 and 212; Alkyl interactions with ALA310, LEU318, and TRP347; sulfur interaction with MET280; Pi-anion interaction with ASP308; and Pi-Pi interactions with HIS 221 and HIS 309. Compound B also had the favorable interactions. However, unlike compound A it had one unfavorable interaction with ALA310 which is due to the donor-donor interaction by the two hydrogen atoms (Fig. 8). The favorable interactions include-hydrogen bonding with ARG 307, GLN 212 and HIS 309; sulfur interaction with HIS 291, Pi-Pi interaction with TRP 347 and alkyl interaction with MET 280 and TYR 282. Like compound B, compound C also has an unfavorable interaction due to the donor-donor interaction between an oxygen atom and Nitrogen atom from compound C and ARG 214, respectively. However, its favorable interactions include-van der Waals interaction with LEU 213; hydrogen bonding interactions with ASP 308, ARG 307 and TRP 347; Pi-sigma interaction with LEU 224; Pi-amide interaction with GLN 212; and alkyl interactions with MET211 and HIS 309. The high number of favorable interactions and absence of unfavorable interaction in compound A is possibly responsible for its highest binding energy (−8.7 kcal/mol) compared to the other selected hits. The presence of GLN 212, HIS 309 and TRP 347 in all the molecular interactions of all three compounds with the protein confirms the same binding pocket.

**Figure 7.**
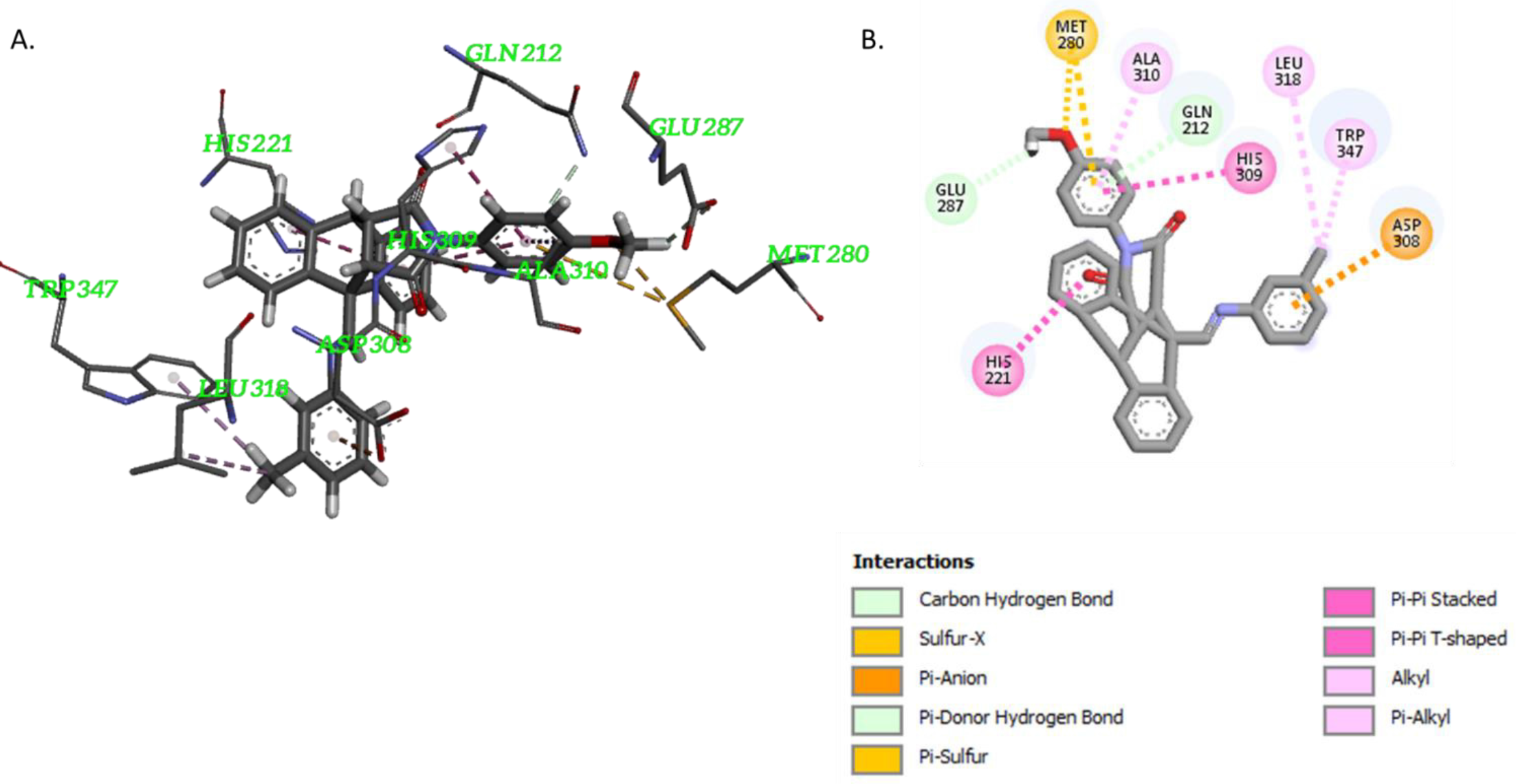
The binding mode (a) and molecular interactions (b) of compound A

**Figure 8.**
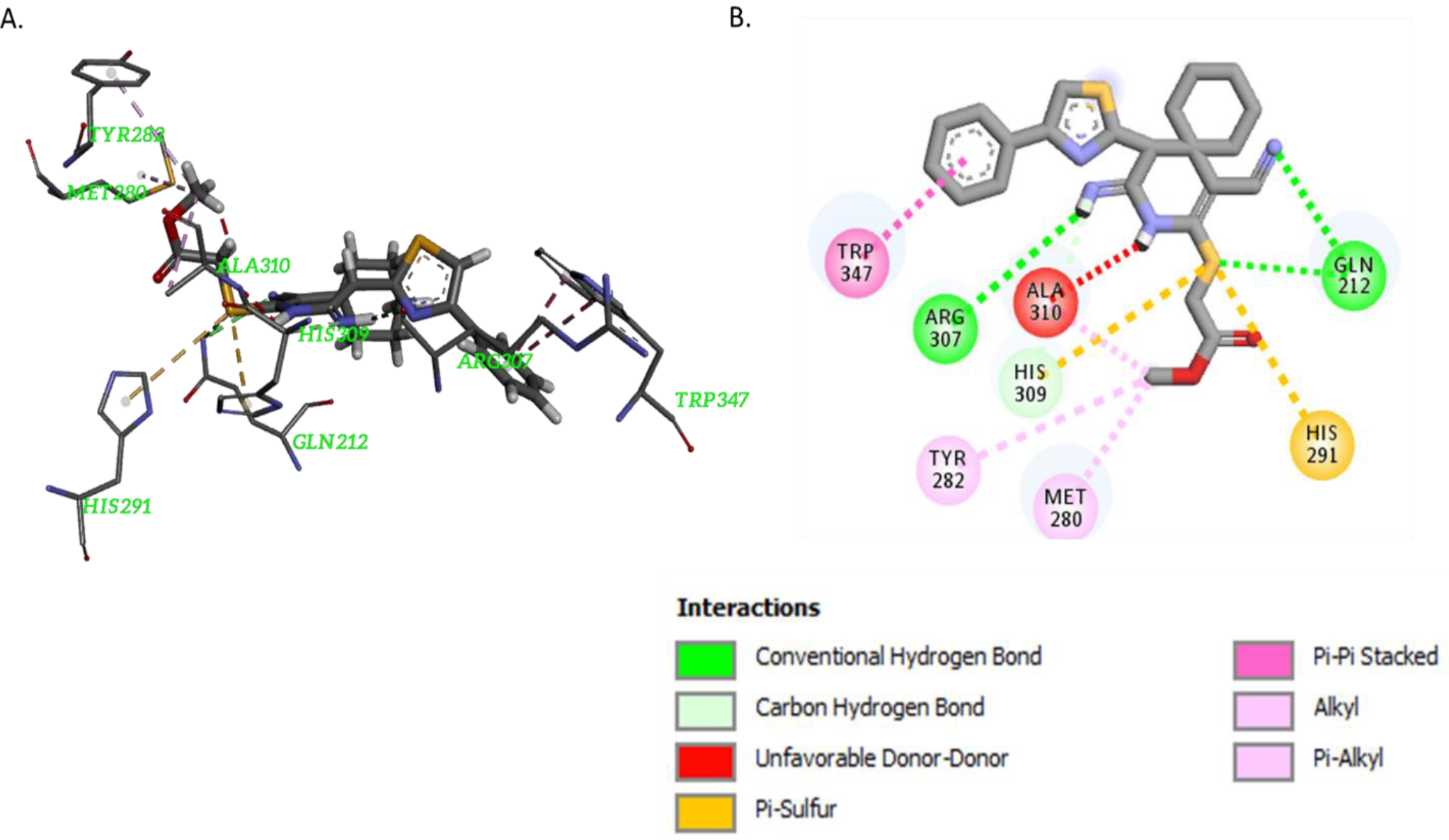
The binding mode (a) and molecular interactions (b) of compound B

**Figure 9.**
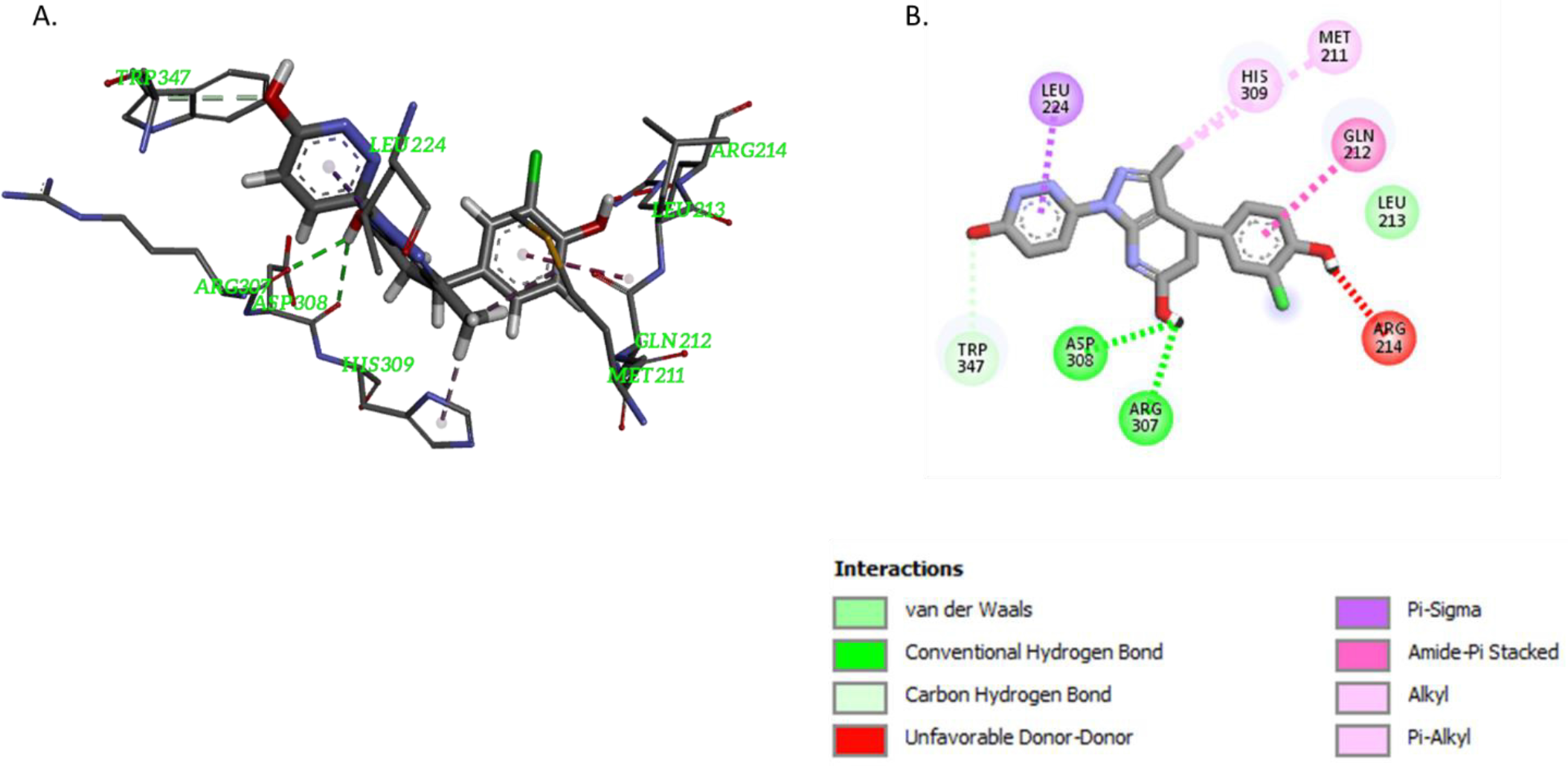
The binding mode (a) and molecular interactions (b) of compound C

### 3.4 Physicochemical and Toxicological Properties of Selected Compounds

The environmental safety of the use of any of the selected hits is quite important as a compound maybe quite effective but due to its toxicological properties maybe quite unsafe. In silico methods have been predominantly used in predicting the physicochemical and toxicological properties in the early stages of discovering potential inhibitors as it is much faster and safer and avoids unnecessary deaths of many animals that may be used for tests (17, 46–50). Moreover, their physicochemical properties (Table 1) which was determined using MCULE (30) is often used for gaining insight into their absorption, distribution and properties. The renowned Lipinski’s RO5 (molar mass and Log P values should not exceed 500 g/mol and 5, respectively; the number of hydrogen bond acceptors (HBD) and hydrogen bond donors (HBA) must not be greater than 10 and 5, respectively) hence, the name RO5 due to the multiples of five being the maximum limit for the rules (32). Only compound C did not violate any of the rules, however, both compounds A and B had Log P values greater than 5. The refractivity, which contributes to the absorption and distribution property of the compound, is expected to be between 40 and 130 (51) of which only compound A exceeded a value of 151.04. Another property, which contributes to the absorption and distribution of a molecule, is the polar surface area (PSA) which is expected to be less than 140 Å^2^ (41) of which compound B exceeded with a value of 152.4. The solubility of the selected hits was predicted using admetsar (52). For a compound to be soluble it is expected to be between −1 and −5 (53) of which all the selected hits qualify. Hence, it can be concluded that only compound C has the most suitable physicochemical properties required among the selected hits.

The toxicity properties (Table 3) were predicted using pkCSM webtool (54), the toxicity checker (30) and DL-AOT (Deep Learning-Acute Oral Toxicity) prediction server (55). The AMES toxicity refers to the mutagenicity of the molecule while the maximum recommended tolerated dose (MRTD) is an estimation of the toxic dose threshold in humans. It is recommended to be less than or equal to 0.477 log mg/kg/day. Fortunately, all the selected hits are safe about mutagenicity and MRTD. The skin sensitization refers to the hazard involved if the compound is applied dermally. The results show that all the selected hits are safe. The LD_50_ (Lethal Dose) for the oral rat acute toxicity considers the toxic potency of a compound. It is the amount of a compound that if given at once would cause the death of 50% of a group of test animals. The DL-AOT webtool predicts the LD_50_ values in four categories namely (1) danger/poison; (2) warning; (3) caution; (4) none required/safe. The values for compounds A, B and C correspond to warning, caution and safe, respectively. The minnow toxicity refers to the LC_50_ (Lethal Concentration) which is the concentration of a compound required to cause the death of 50% of a group of flathead minnows. A value below 0.5 log mM is regarded as high acute toxicity. Only compound C was predicted to be safe with a value of 2.014. The toxicity checker (30) tool was used to deduce the toxicity alerts in the selected hits. However, it depicted that the N-C=O group in the imidazole ring and the methoxy group as the toxic alerts for compound A while the ester group as the toxic alert for compound B. Nevertheless, it showed no alerts for compound C confirming it as a safe compound.

**Table 2.**
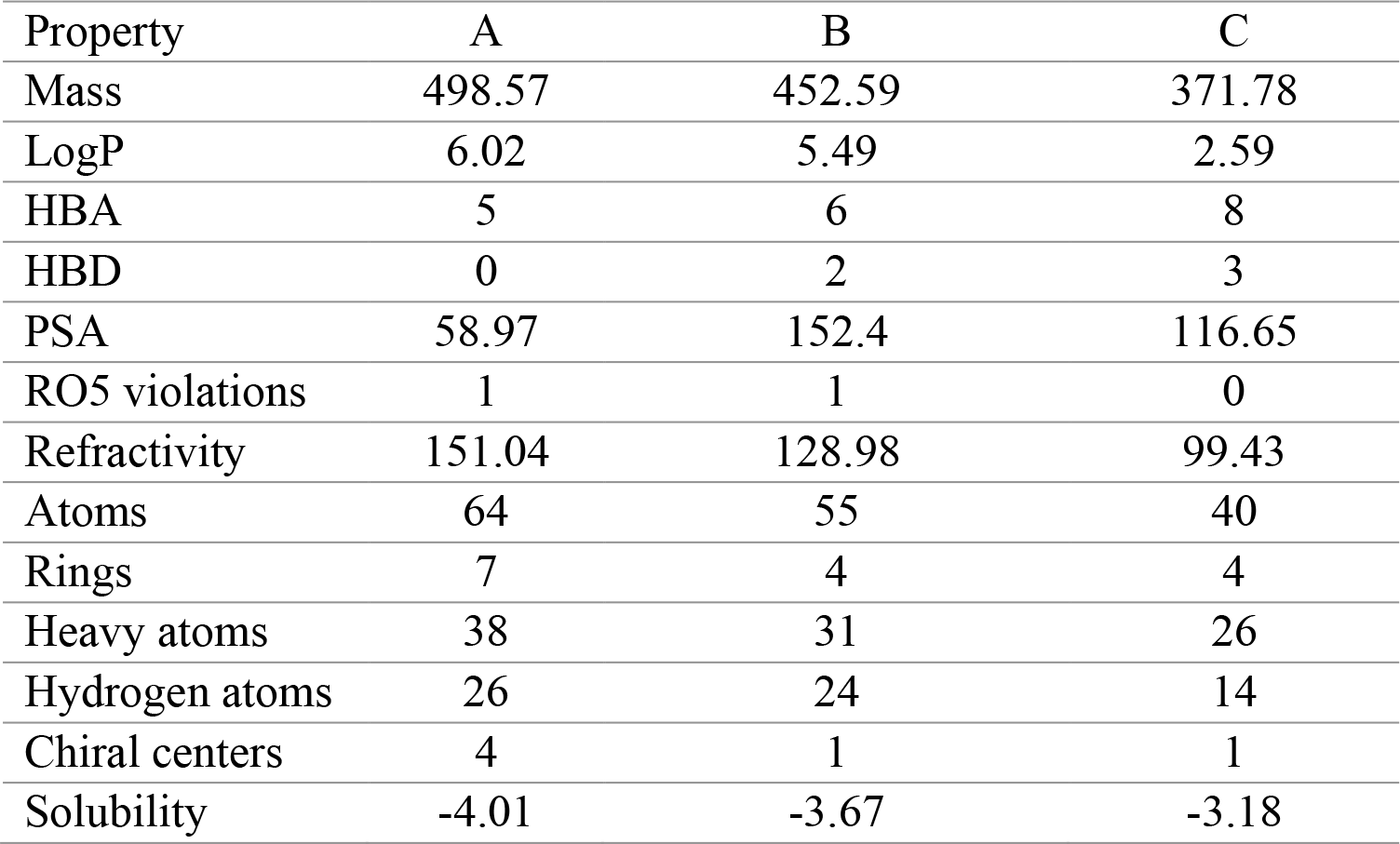
The Physicochemical properties of the selected HITS

| Property | A | B | C |
| --- | --- | --- | --- |
| Mass | 498.57 | 452.59 | 371.78 |
| LogP | 6.02 | 5.49 | 2.59 |
| HBA | 5 | 6 | 8 |
| HBD | 0 | 2 | 3 |
| PSA | 58.97 | 152.4 | 116.65 |
| RO5 violations | 1 | 1 | 0 |
| Refractivity | 151.04 | 128.98 | 99.43 |
| Atoms | 64 | 55 | 40 |
| Rings | 7 | 4 | 4 |
| Heavy atoms | 38 | 31 | 26 |
| Hydrogen atoms | 26 | 24 | 14 |
| Chiral centers | 4 | 1 | 1 |
| Solubility | -4.01 | -3.67 | -3.18 |

**Table 3.** The toxicological properties of the selected hits

| Properties | A | B | C | Units |
| --- | --- | --- | --- | --- |
| AMES toxicity | No | No | No | Categorical (Yes/No) |
| Max. tolerated dose (human) | 0.302 | -0.573 | -0.151 | Numeric (log mg/kg/day) |
| Skin Sensitization | No | No | No | Categorical (Yes/No) |
| Oral Rat Acute Toxicity (LD50) | 2.77 | 2.97 | 3.16 | log(mg/kg)) |
| Minnow toxicity | -2.262 | -0.987 | 2.014 | Numeric (log mM) |

**Figure 10.**
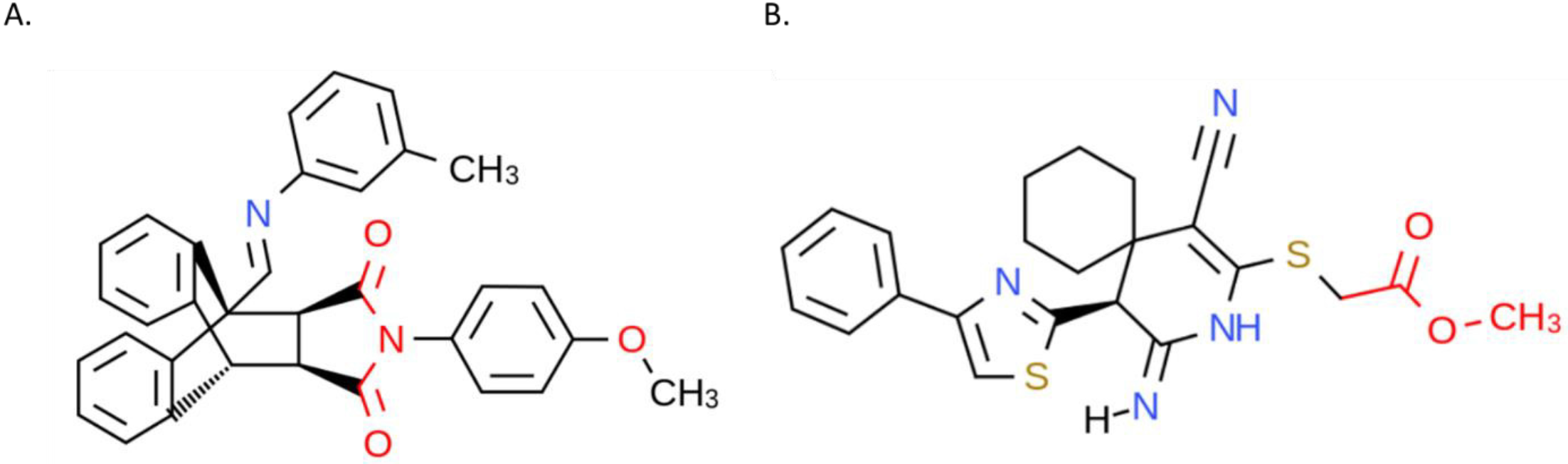
The toxicity alerts in red font for compounds A and B

## 4. Conclusions

The use of virtual screening and applying a consensus scoring method, which combines both molecular docking and machine learning, helped in selecting three compounds as hits from a database of about 40 million compounds. The molecular docking results showed that all the compounds have negative binding energy with compound A having the highest docking score. The molecular interactions revealed that the high binding affinity observed in compound A is most likely due to the high number of favorable interactions and absence of unfavorable interactions, which occurred compounds B and C. However, based on the physicochemical and toxicological properties. Compound C is the best choice, as it does not violate any of the Lipinski’s RO5, and other recommended properties that relate to absorption and distribution properties. Moreover, compound C was predicted to be the only safe compound with respect to LD_50_ and LC_50_ for both the acute toxicity for both rats and flathead minnows. Nevertheless, compound C can be modified to increase its binding affinity by replacing the hydroxyl group, which is a hydrogen bond donor and is responsible for the unfavorable interaction, with a carbonyl group which is a hydrogen bond acceptor and hence would form a favorable interaction with ARG 214.. Nevertheless, whatever modification(s) are made to compound C it is important to take into consideration the effect it would have on its physicochemical and toxicological properties. Consequently, compound A could be modified using the toxic alerts as a guide. The oxygen in the methoxy group could be replaced with a nitrogen atom making it a tertiary amine. Also, the two carbonyl groups attached to the imidazole ring should be replaced with any halogen atom. Summarily, when modifying selected hit compounds, there is often a tradeoff between increasing binding affinity and acquiring good physicochemical and toxicological properties. It is recommended that compound C should be validated experimentally to confirm its inhibitory activity against *Desulfobacter postgatei* (a prominent sulfate reducing bacteria). A positive result would reduce the level of sulfur produced in oil and gas wells, which will subsequently prevent formation of iron sulfide scales such as pyrite. These findings will help in directing future experimental research on this subject.

## Acknowledgment

This publication was made possible by NPRP Grant # 9-084-2-041 from Qatar National Research Fund (a member of Qatar Foundation). The findings achieved herein are solely the responsibility of the authors. Qatar University and the Gas Processing Center are acknowledged for their support.

